# Spermidine facilitates the adhesion and subsequent invasion of *Salmonella* Typhimurium into epithelial cells via the regulation of surface adhesive structures and the SPI-1

**DOI:** 10.1101/2023.06.03.543567

**Authors:** Abhilash Vijay Nair, Utpal Shashikant Tatu, Yashas Devasurmutt, S.A Rahman, Dipshikha Chakravortty

## Abstract

Polyamines are poly-cationic molecules ubiquitously present in all organisms. *Salmonella* synthesizes and also harbors specialized ABC transporters to uptake polyamines. Polyamines assist in pathogenesis and stress resistance in *Salmonella*; however, the mechanism remains elusive. The virulence trait of *Salmonella* depends on the injection of effector proteins into the host cell and modulation of host machinery and employs an array of arsenals to colonize in the host niche successfully. However, prior to this, *Salmonella* utilizes multiple surface structures to attach and adhere to the surface of the target cells. Our study solves the enigma of how polyamine spermidine assists in the pathogenesis of Salmonella. We show that spermidine mediates the initial attachment and adhesion of *Salmonella* Typhimurium to Caco-2 cells, facilitating its invasion. In-vivo studies showed that polyamines are required for invasion into the murine Peyer’s patches. Polyamines have previously been shown to regulate the transcription of multiple genes in both eukaryotes and prokaryotes. We show that spermidine controls the RNA expression of the two-component system, BarA/SirA, that further regulates multiple fimbrial and non-fimbrial adhesins in *Salmonella*. Flagella is also a vital surface structure aiding in motility and attachment to surfaces of host cells and gall stones. Spermidine regulated the expression of flagellin genes by enhancing the translation of s28, which features an unusual start codon and a poor Shine-Dalgarno sequence. Besides regulating the formation of the adhesive structures, spermidine tunes the expression of the *Salmonella* pathogenicity island-1 encoded genes. Thus, our study unravels a novel mechanism by which spermidine aids in the adhesion and the subsequent invasion of *Salmonella* into host cells.

## Introduction

*Salmonella enterica* is considered a primary foodborne pathogen and the most pathogenic species of the genus *Salmonella* [1]. This species comprises more than 2500 serovars broadly classified into Typhoidal (TS) and Non-typhoidal forms (NTS). The serovars *S.* Typhimurium and *S.* Eneteridis infect a wide range of hosts leading to diarrhoea and gastroenteritis, while the human-restricted serovars, *S.* Typhi and *S.* Paratyphi cause severe systemic infection and enteric fever [2, 3]. However, in children and immunocompromised individuals, the NTS can often cause systemic disease and fever-like symptoms classified as invasive non-Typhoidal serovars (iNTS) [4]. Upon ingestion through contaminated food and water, *Salmonella* survives the acidic pH in the stomach and successfully reaches the small intestine. *Salmonella* harbours multiple virulence-associated genes, most of which are clustered into 23 *Salmonella* pathogenicity islands (SPIs) [5]. The primary cell type encountered by *Salmonella* is the epithelial cells lining the intestinal lumen at the Peyer’s patches. The first step to a successful infection is passing through the thick mucous lining and adhering to the epithelial cell surface followed by its subsequent invasion[6]. To infect the host cells at the Peyer’s patches, it utilises a highly elegant nanomachine called the Type 3 secretion system (T3SS) encoded by the SPI-1, which transports effectors into the host cytosol leading to actin cytoskeletal rearrangement and uptake of the bacteria [7]. Once inside the host cell, it activates another set of virulence genes encoded by SPI-2, which aid in the survival and replication of *Salmonella* in the specialised niche called the *Salmonella* containing vacuole (SCV) [7, 8]. It is then taken up by the CD18+ expressing macrophages and dendritic cells and disseminates through the reticuloendothelial system [9, 10].

Polyamines are positively charged compounds that are associated with a number of functions in eukaryotes including cell growth, cell division, stress response gene regulation etc [11]. They are ubiquitously present in all life forms. *Salmonella* also has the ability to utilise polyamines. *Salmonella* can anabolically metabolise putrescine, spermidine and spermine from arginine and ornithine as the precursor[11]. Besides its ability to synthesise, it has three transport systems: PotABCD, PotFGHI and PotE. PotABCD imports mainly spermidine and putrescine, while PotFGHI PotE imports and exports only putrescine respectively [12].

In bacteria, polyamines are involved in biofilm formation, virulence, and motility [13, 14]. In *Shigella sp.* the accumulation of spermidine shields it from oxidative stress within the host macrophages, which is essential for its invasion [15]. During infection of *Streptococcus pneumoniae*, spermidine transporter PotD levels increase and PotD loss leads to attenuated pneumococcal virulence in mice [16]. Also, in *Vibrio cholerae* and *Yersinia pestis,* polyamines are crucial players regulating biofilm formation. In *Salmonella* Typhimurium polyamines are crucial for the virulence and stress resistance [17]. *C. elegans* infected with *Salmonella* lacking the ability to synthesise both putrescine and spermidine and with *Salmonella* unable to transport the two polyamines survive better than infected with the wild type [12]. Also, the deletion of *speG* that functions in the catabolism of spermidine in *Salmonella* Typhimurium reduces the intracellular replication in multiple human cell lines [18]. Spermidine is a major polyamine in bacteria and functions to regulate the expression of numerous genes in prokaryotes by interacting directly with the negatively charged nucleic acids. Spermidine and putrescine are the two vital polyamines present in *Salmonella*. All the studies till date have shown the function of a group of polyamines in *Salmonella.* However the mechanism involved in assisting in virulence of this bacteria by polyamines still remains a mystery.

This led us to ask the question of whether spermidine alone plays a role in the pathogenesis of *Salmonella* Typhimurium. Also, to unravel the mechanism by which spermidine modulates the virulence of *Salmonella* Typhimurium. In our study, we observe that spermidine is an essential player during the infection cycle of *Salmonella* Typhimurium. Spermidine is critical in each of the early stages of the infection cycle of *Salmonella* and its survival within host epithelial cells. We delineate a novel regulatory network involving spermidine in modulating the bacterial surface adhesive and motility structures in *Salmonella* Typhimurium.

## Material and Methods

### Bacterial strains and growth condition

*Salmonella enterica* serovars Typhimurium (STM WT) wild type strain ATCC 14028 was used in all experiments which was a kind gift from Prof. Michael Hensel, Abteilung Mikrobiologie, Universität Osnabrück, 273 Osnabrück, Germany. The bacterial strain was cultured in Luria broth (LB-Himedia) with constant shaking (175rpm) at 37°C orbital-shaker. Kanamycin, Chloramphenicol and Ampicillin antibiotics (Kanamycin-50μg/ml, Cholramphenicol-20μg/ml and Ampicillin-50μg/ml) were used wherever required. For immunofluorescence studies, the strains were transformed with pFPV 25.1 plasmid expressing mCherry.

### Bacterial gene knock-out and strain generation

The generation of gene knock-out in bacteria was done using the One-step chromosomal gene inactivation protocol by Datsenko and Wanner (2000) [19]. Briefly, primers were designed for amplification of Kanamycin resistance gene and Chloramphenicol resistance gene cassettes from PKD4 and PKD3 plasmids, respectively, such that 5’terminus of the primers had sequence homologous to the flanking region of the gene to be knocked out (here *potCD and speED*). After amplifying the Kanamycin resistance gene and the Chloramphenicol resistance gene, the amplified products were purified using chloroform-ethanol precipitation. Followed by electroporation into the STM WT cells (expressing PKD-46 plasmid which provides the λ-Red recombinase system) by a single pulse of 2.25 kV separately for the Kan^R^ and Chlm^R^. Immediately fresh recovery media was added and incubated at 37°C for 60 minutes in an orbital-shaker. After incubation, the cultures were centrifuged at 10000rpm for 5 minutes. Then the pellet was dissolved in 100μL of media and plated at the required dilution on LB agar plates with Kanamycin and Cholaramphenicol as required. The plates were incubated at 37°C for 12-14 hours. The colonies were selected and patched on fresh plates and confirmed for knock-out using PCR with primers designed corresponding to the ∼100bp upstream and downstream of the genes (knocked out) for both the knock-out strains. Which was then run on 1% agarose gel to compare the length of the products in the mutants from STM WT bacteria. For the generation of the double knock-out strain (STM *ΔpotCDΔspeED*), the STM *ΔpotCD* (resistant to Kanamycin) was first transformed with the plasmid pKD46 which provides the λ-Red recombinase system. To this transformed strain, the purified PCR product to knock-out *speED* was electroporated to generate the STM *ΔpotCDΔspeED* (resistant to Kanamycin and Chloramphenicol). For generation of chromosomal *fliA*-FLAG in STM WT, STM *ΔpotCD,* and STM *ΔspeED* the same protocol of homologous recombination using pKD-46 plasmid was used. The amplification of FLAG-Kan^R^ was done from pSUB11 plasmid (a generously gift from Prof. Umesh Varshney, MCB, IISc). The purified PCR product (FLAG-Kan^R^ flanked by homologous sequence for incorporation at the 3’-end of *fliA* excluding the STOP codon) were electroporated into the respective strains and transformants were selected in Kanamycin containing LB agar plates.

### Cell culture and maintenance

Caco-2 cells (human intestinal epithelial cell line) were cultured in DMEM - Dulbecco’s Modified Eagle Medium (Lonza) supplemented with 10% FBS (Gibco), 1% Non-essential amino acids (Sigma-Aldrich), 1% Sodium pyruvate (Sigma-Aldrich) and 1% Penicillin-streptomycin (Sigma-Aldrich) at 37°C humidified chamber (Panasonic) with 5% CO_2_. HeLa cells (human epithelial cell line) were cultured in DMEM - Dulbecco’s Modified Eagle Medium (Lonza) supplemented with 10% FBS (Gibco) at 37°C humidified chamber (Panasonic) with 5% CO_2_. For each experiment, the cells were seeded onto the appropriate treated cell culture well plate at a confluency of 80% either without coverslips (for intracellular survival assay, adhesion assay and qRT-PCR) or with coverslips (for immunofluorescence microscopy).

### Gentamicin protection assay

The cells were infected with STM WT, STM *ΔpotCD,* STM *ΔspeED* and STM *ΔpotCDΔspeED* at MOI of 10 (for intracellular survival assay, adhesion assay, and immunofluorescence microscopy) and MOI 25 (for qRT-PCR). After infecting the cell line with STM WT and the mutants, the plate was centrifuged at 700-900 rpm for 10 minutes to facilitate the proper adhesion. The plate was then incubated for 25 minutes at 37°C humidified chamber and 5% CO_2_. Then the media was removed from the wells and washed with 1X PBS. Fresh media containing 100 µg/mL gentamicin was added and again incubated for 60 minutes at 37°C and 5% CO_2_. The media was then removed, cells were washed with 1X PBS twice, and fresh media containing 25µg/mL gentamicin was added. The plate was incubated at 37°C and 5% CO_2_ till the appropriate time. For the intracellular survival assay, two time points were considered 2 hours and 16 hours, and for qRT-PCR three time points were considered 2 hours, 6 hours and 16 hours.

### Intracellular survival assay and invasion assay

At the appropriate time post-infection the cells were lysed using 0.1% Triton X followed by addition of more 1X PBS and samples were collected. The collected samples were plated at the required dilutions on LB agar plates and kept at 37°C. 12 hours post incubation the Colony forming units (CFU) were enumerated for each plate.

The fold proliferation and invasion were determined as follows

Fold Proliferation = (CFU at 16 hours post-infection)/(CFU at 2 hours post-infection)

Percentage invasion = [(CFU at 2 hours post-infection)/(CFU of the Pre-inoculum)]×100

### Adhesion assay

The cells were infected with STM WT, STM *ΔpotCD,* STM *ΔspeED* and STM *ΔpotCDΔspeED* at MOI of 10. After infection, the plate was centrifuged at 700-900 rpm for 10 minutes to facilitate the proper adhesion. The plate was then incubated for 10 minutes at 37°C humidified chamber and 5% CO_2_. Then the media was removed, and the cells were washed with 1X PBS twice to remove the loosely adhered bacteria. The mammalian cells were then lysed with 0.1% Triton-X 100 to release the adhered bacteria into the solution, and more of 1XPBS was added and the samples were collected. The collected samples were plated at the required dilutions on LB agar plates and kept at 37°C. 12 hours post incubation, the Colony forming units (CFU) were enumerated for each plate. The percentage adhesion was determined as follows:

Percentage adhesion = [(CFU at 10 minutes post-infection)/ (CFU of the Pre-inoculum)]×100

### Confocal microscopy

After the appropriate incubation time, the media was removed, and the cells were washed with 1X PBS. The cells were then fixed with 3.5% Paraformaldehyde for 10 minutes. The cells were then washed with 1X PBS and incubated with the required primary antibody (anti-*Salmonella* LPS) in a buffer containing 0.01% saponin and 2% BSA and incubated at room temperature for 45-60 minutes. The primary antibody was then removed by washing with 1X PBS and then incubated with the appropriate secondary antibody tagged to a fluorochrome. The coverslips were then washed with PBS and mounted on a clean glass slide using mounting media containing an anti-fade reagent. The coverslips were sealed with clear nail polish and observed under the confocal microscope (Zeiss 710 microscope, at 63X oil immersion, 2×319 3x zoom, and 100X zoom for studying only bacterial samples, Zeiss 880 microscope, at 63X oil immersion, 2×319 3x zoom). For studying histopathology samples 40X oil immersion 2×319 3x zoom was used. For FliC study, STM WT, STM *ΔpotCD,* STM *ΔspeED* were subcultured and grown in LB media, with or without supplementation of 100µM spermidine till log phase of growth (OD 0.1). After washing with 1X PBS, the samples were smeared on clean glass slide and air dried. Followed by staining as previously explained using specific antibody in buffer. For invasion assay buffer contained only 2% BSA.

### RNA isolation and qRT-PCR

RNA isolation was performed from infected cells after appropriate hours of infection with STM WT, STM *ΔpotCD,* STM *ΔspeED* by RNA isolation was performed using TRIzol (Takara) reagent according to manufactures’ protocol RNA was quantified using Thermo-fischer scientific Nano Drop followed by running on 2% agarose gel for checking the quality. For cDNA synthesis, first DNase I treatment with 3μg of isolated RNA was done at 37℃ for 60 minutes, which was then stopped by heating at 65℃ for 10 minutes. Then RNA (free of DNA) was subjected to Reverse transcription using Random hexamer, 5X RT buffer, RT enzyme, dNTPs and DEPC treated water at 37°C for 15 minutes, followed by heating at 85℃ for 15 seconds. Quantitative real-time PCR was done using SYBR green RT-PCR kit in BioRad qRT-PCR system. A 384 well plate with three replicates for each sample was used. The expression levels of the gene of interest were measured using specific RT primers. Gene expression levels were normalised to 16SrDNA primers of *S*. Typhimurium. For expression studies in bacteria grown in LB media, the bacterial samples were harvested at 3 hours, 6 hours, 9 hours and 12 hours post subculture in fresh LB media in 1:100 ratio. Then similar protocol was used to isolate total RNA using TRIzol (Takara) reagent according to manufactures’ protocol.

### Immunoblotting

STM WT, STM *ΔpotCD,* STM *ΔspeED* and STM *ΔpotCDΔspeED* were grown in LB media until log phase of growth. The cells were centrifuged to remove the media and the cells were resuspended in lysis buffer (Sodium chloride, Tris, EDTA, 10% protease inhibitor cocktail) after washing with 1XPBS. The cells were lysed using sonication and centrifuged at 4°C to collect the cell lysate, followed by estimation of total protein using the Bradford protein estimation method. 50µg of protein was loaded on to a Polyacrylamide Gel Electrophoresis (PAGE), then transferred onto 0.45μm PVDF membrane (GE Healthcare). 5% skimmed milk (Hi-Media) in TTBS was used to block for 1h at RT and then probed with Anti-FLAG primary and secondary HRP-conjugated antibodies. ECL (Biorad) was used for developing the blot, and images were captured using Chemi-Doc GE healthcare. All densitometric analysis was performed using the Image J.

### Swim assay

2 µl of bacterial samples were spotted on the 0.3% agar plates supplemented with 0.5% yeast extract, 1% casein enzyme hydrolysate, 0.5% NaCl and 0.5% glucose (swim agar plates). The plates were incubated at 37°C for 6 hours, and then images were taken using a Biorad-chemidoc. The diameters of the motility halos were measured using ImageJ. At least five replicate plates were used for each condition.

### Transmission electron microscopy

Flagella were visualised by slightly modifying the protocol described in Garai et al 2016 [20]. Briefly, overnight STM WT, STM *ΔpotCD,* STM *ΔspeED* and STM *ΔpotCDΔspeED* were inoculated in LB media(1:33 ratio) and incubated at 37°C orbital shaker incubator for 2-3 hours until it reached an OD of 0.2. The bacterial cultures were centrifuged at 2000rpm for 10 minutes at 4°C. The bacterial cells were washed with 1XPBS twice and finally the cells were resuspended in 100 µl of 1X PBS. Similarly, STM *ΔfliC* was used as negative control. 10 µl of the cell suspension was added on copper grid, air dried, stained with 1% uranyl acetate for 30 sec, and visualised under transmission electron microscope.

### Mass-spectrometry for determination of intracellular spermidine

The sample preparation was done as explained Feng Y et.al; previously[21]. Briefly, STM WT, STM *ΔpotCD,*STM *ΔspeED* and STM *ΔpotCDΔspeED* were grown in LB media until log phase of growth. The cells were centrifuged to remove the media and the cells were resuspended in 80% methanol (Thermo-fischer) after washing with 1XPBS. The cells were lysed using sonication and centrifuged at 4°C to collect the cell lysate. The methanol was evaporated in vacuum at low temperatures, and then lyophilised at -40°C. Dried metabolite extracts were dissolved in 1.0 mL of 0.1% Formic acid in ddH2O and vortexed for 2 minutes, later centrifuged for 5.0 minutes at 5000 rpm and 4°C. 0.5 ml of supernatant was transferred to a HPLC vial (Amber colored) for LC–MS/MS analysis on a Agilent 1260 HPLC system coupled to an Agilent QQQ 6460 mass spectrometer. An Agilent Eclipse Plus C18 (50 mm × 4.6 mm, 1.8 μm) column at 30°C was utilized for LC separation. Samples were injected (10 μl) from the auto sampler kept at 5°C. Then mobile phase A (ddH2O containing 0.1% formic acid) and mobile phase B (Methanol containing 0.1% formic acid) were prepared for sample elution. The isocratic elution was as follows: 20% mobile phase B was maintained for 5 min at a flow rate of 0.5 ml min−1. Mass spectra were acquired on a 6460 QQQ mass spectrometer (Agilent, USA) equipped with an electrospray ionization (ESI) source in positive ion mode. The concentrations 1, 5, 10, 25, 50, 75, 100 ng/ml of standard (fresh preparations) were used for Linearity. The Mass parameters and multiple reaction monitoring transition ions of spermidine are shown in Supplementary Table. Peak identification and amounts of spermidine were evaluated using Agilent MassHunter Data Acquisition, Agilent MassHunter QQQ Qualitative Analysis and Agilent MassHunter QQQ Quantitative Analysis softwares on the basis of the known amounts of spermidine.

### In-vivo animal experiment

5-6weeks old C57BL/6 mice were infected by orally gavaging 10^7^ CFU of STM WT, STM *ΔpotCD,* STM *ΔspeED* and STM *ΔpotCDΔspeED*. For invasion assay intestine was isolated 6 hours post-infection, and CFU was enumerated on differential and selective SS agar by serial.

### Availability of data and materials

All data generated and analysed during this study, including the supplementary information files, are incorporated in this article. The data is available from the corresponding author on request.

### Ethics statement

All the animal experiments were approved by the Institutional Animal Ethics Committee, and the Guidelines provided by National Animal Care were strictly followed during all experiments. (Registration No: 435 48/1999/CPCSEA).

## Results

### Spermidine transporter and biosynthesis genes are co-regulated in *Salmonella* Typhimurium

*Salmonella* Typhimurium harbours both the ability to synthesise spermidine and import it from the extracellular milieu. This intrigued us to investigate the regulation of spermidine synthesis and its transport by *Salmonella.* To begin with we accessed the expression of the genes *potA*, *potB*, *potC*, and *potD,* which encode the ATP-dependent transport apparatus that localises to the cell wall passing through the periplasmic space [22]. We observed that during *in-vitro* growth of *Salmonella* in LB media, all the genes encoding the transporter show a bimodal mRNA expression, with 8-fold, 6-fold, 11-fold and 9-fold higher expression at the mid-log phase (6 hours) than the early-log phase(3 hours) for *potA*, *potB*, *potC* and *potD* respectively. Followed by a dip in the expression at the late log phase(9 hours), *potA*, *potB*, *potC,* and *potD* show a 6-fold, 3-fold, 7-fold and 5-fold upregulation in the mRNA expression, respectively, at the early stationary phase(12 hours) (**Fig 1 A**). Similarly, we observed that *speE* and *speD,* the genes encoding the two enzymes that catalyse the synthesis of spermidine and decarboxylation of S-Adenosyl methionine, show a 7-fold and 8-fold upregulation in the mRNA expression at the mid-log phase and 5-fold and 7-fold upregulation in the mRNA expression at the early stationary phase of *in*-*vitro* growth in *Salmonella* Typhimurium (**Fig 1 B**). In prokaryotes, spermidine plays a pivotal role in growth, stress response, nutrient starvation, etc., which explains the higher mRNA expression of the transport and biosynthesis genes during the mid-log phase in *Salmonella*. However, a question remains: how does *Salmonella* regulate the transport system and its intracellular spermidine biosynthesis during its growth? To solve this mystery, we generated chromosomal knock-out strains of *Salmonella* Typhimurium, namely STM *ΔpotCD* that cannot import spermidine, STM *ΔspeED* that cannot synthesise spermidine and a double knock-out STM *ΔpotCDΔspeED* that lacks both the functions. We studied the mRNA expression of biosynthesis genes in transporter mutants and vice-versa. We observed that both *speE* and *speD* show downregulation post the early-log phase in STM *ΔpotCD* compared to their mRNA expression in STM WT during their *in-vitro* growth in LB media (**Fig- 1 C**). Likewise, *potA*, *potB*, *potC* and *potD* showed significant downregulation in mRNA expression post the early-log phase compared to their mRNA expression in STM *ΔspeED* compared to their mRNA expression in STM WT (**Fig 1 D**). Thus in *Salmonella,* spermidine import and biosynthesis are co-regulated. To further verify our observation, we determined the intracellular levels of spermidine in the different strains, and we observed that, indeed, the spermidine levels in both the mutants STM *ΔpotCD* and STM *ΔspeED* were significantly lower than STM WT (**Fig 1 E, Fig S1 D and E**). These results indicate that in *Salmonella* Typhimurium, spermidine import, and biosynthesis are higher during the mid-log phase, and both of these functions are co-regulated and are not mutually exclusive. Further, we analysed how the loss of spermidine import and biosynthesis affects the growth of *Salmonella* in rich media such as LB medium, minimal media such as M9 media and F-media that mimics the acidic environment of SCV. In LB media, all the strains showed similar growth kinetics (**Fig S1 A**). Similarly, we did not observe any difference in minimal M9 media and acidic F-media (**Fig 1 F, Fig S1 B-D**). We further supplemented spermidine and putrescine in minimal M9 media during the *in-vitro* growth; however, we also did not observe any significant difference upon supplementation (**Fig. 1 G and H**). Thus, loss of spermidine import and biosynthesis does not alter the growth kinetics of *Salmonella in-vitro*.

**Fig 1:**
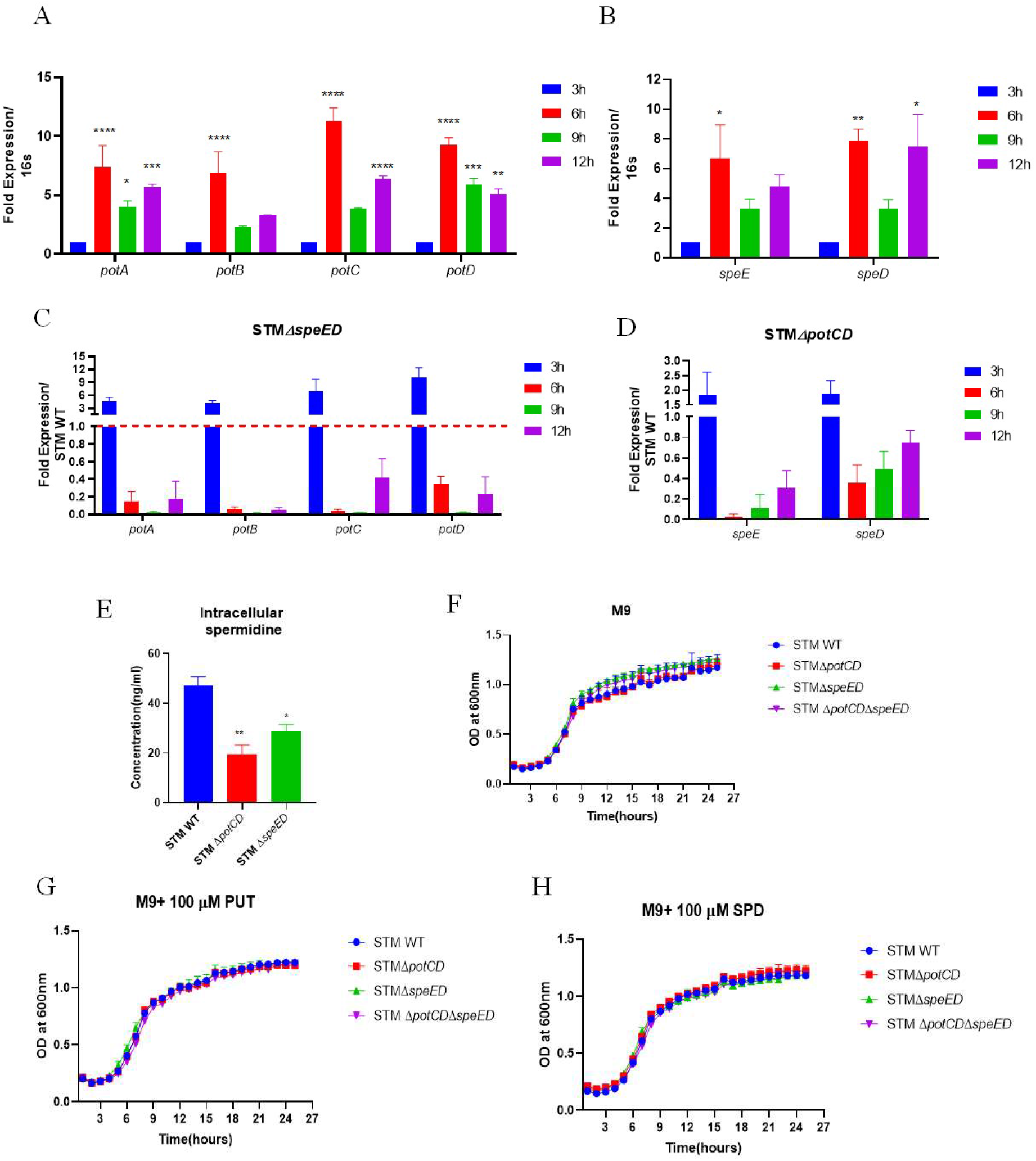
Spermidine transporter and biosynthesis genes are co-regulated in *Salmonella* Typhimurium. . A. The mRNA expression of *pot-*transporter genes in STM WT during *in-*vitro growth in LB media. B. The mRNA expression of *speE* and *speD* genes in STM WT during *in-*vitro growth in LB media C. The mRNA expression of *pot-*transporter genes in STM Δ*speED* during *in-* vitro growth in LB media D. The mRNA expression of *speE* and *speD* genes in STM Δ*potCD* during *in-*vitro growth in LB media E. Intracellular spermidine determination using Mass spectrometry F. Growth kinetics of STM WT, STM Δ*potCD,* STM Δ*speED* and STM Δ*potCD* Δ*speED* in M9 minimal media, G. in M9 minimal media supplemented with 100µM Putrescine (PUT), H. in M9 minimal media supplemented with 100µM Spermidine (SPD). Student’s t-test was used to analyze the data; p values ****<0.0001, ***<0.001, **<0.01, *<0.05.

### Spermidine synthesis and transport in *Salmonella* Typhimurium is essential to invade and proliferate in the host

*Salmonella* infects the host system through the faecal-oral route via contaminated food and water [23]. The primary site of infection is the intestinal epithelial cells (IECs) in the large intestine [24]. Thus, we explored the ability of spermidine mutants to invade followed by proliferate into human intestinal cell line Caco-2 cells. We infected Caco-2 cells with all the strains and determined the percentage invasion and the fold proliferation. We observed that STM *ΔpotCD*, STM *ΔspeED*, and STM *ΔpotCD ΔspeED* invaded significantly less than STM WT (**Fig 2 A**). Also, all the mutants exhibited a substantially lower fold proliferation than STM WT in Caco-2 cells (**Fig 2 B**). To validate these results, we performed the infection into HeLa cells, and likewise, we observed a similar lesser invasion and lower fold proliferation of the spermidine transport and biosynthesis gene mutants (**Fig S2 A and B**). The crucial step during the infection of *Salmonella* is its ability to invade the IECs successfully [25, 26]. Upon invasion into these nonphagocytic IECs, the bacteria reside in the SCV and, employing multiple arsenals, survive and proliferate [27, 28]. The above results led us to study the expression of the transporters and the biosynthesis genes of STM upon infection into Caco-2 cells. The transporters *potA*, *potB*, *potC* and *potD* showed a gradual upregulation of their corresponding mRNA expression post 2 hours of infection into Caco-2 cells (**Fig 2 D**). Furthermore, the *speE* and *speD* showed a gradual upregulation of mRNA expression post 2 hours of infection in Caco-2 cells (**Fig 2 C**). All the genes showed a significant upregulation at 16 hours post-infection (**Fig 2 C and D**). These results might explain the lower fold proliferation in Caco-2 cells upon deletion of the transporter and biosynthesis genes in *Salmonella* Typhimurium. As previously studied, we further determined the regulation of the two sets of genes during infection into Caco-2 cells. We observed a similar downregulation of *speE* and *speD* mRNA expression in STM *ΔpotCD* and downregulation of *potA*, *potB*, *potC* and *potD* mRNA expression in STM *ΔspeED* upon infection into Caco-2 cells (**Fig 2 E and F**). These results explain the lower fold proliferation of both the individual mutants even though they lack just one of the two functions (transport/ biosynthesis).

**Fig 2:**
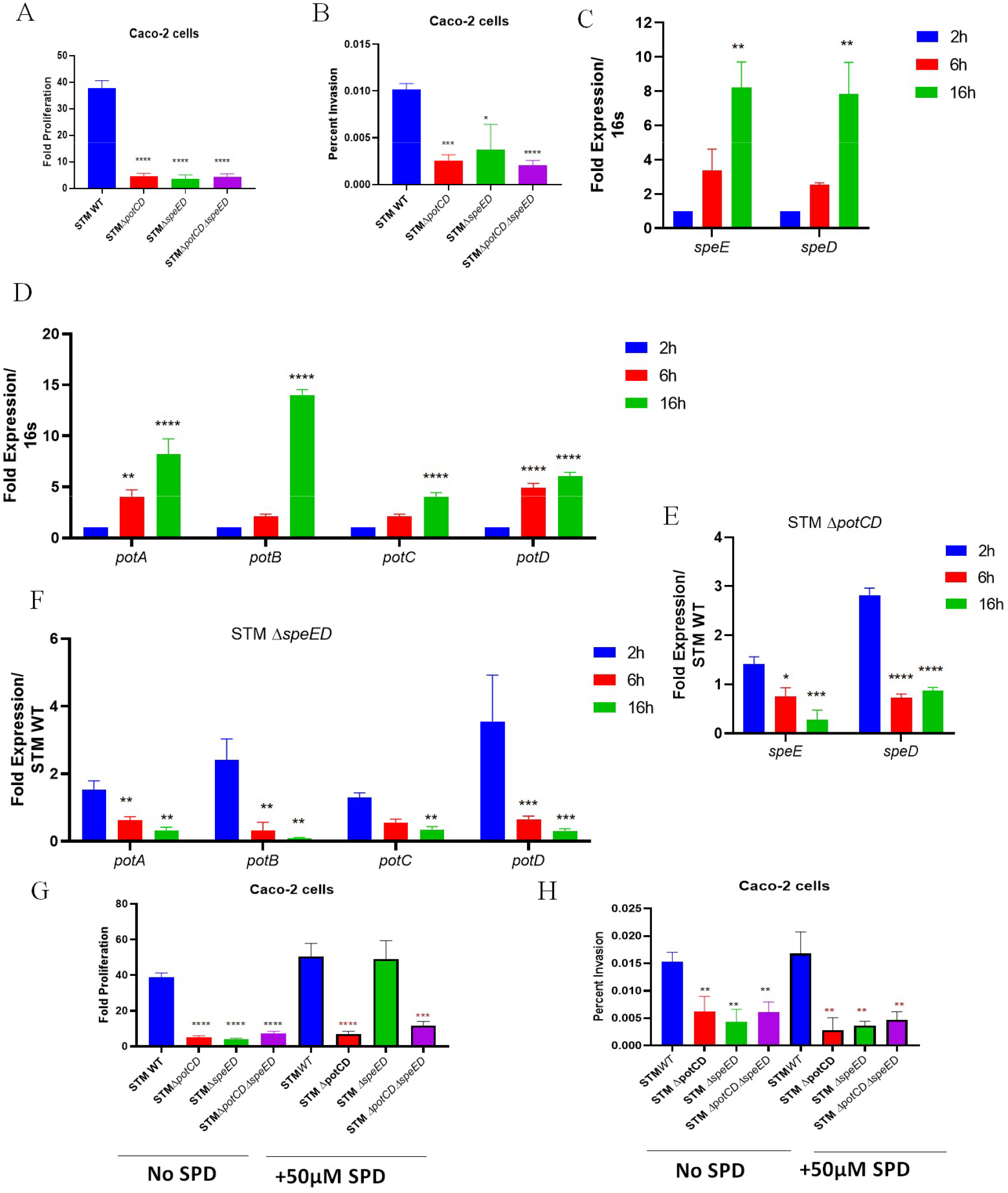
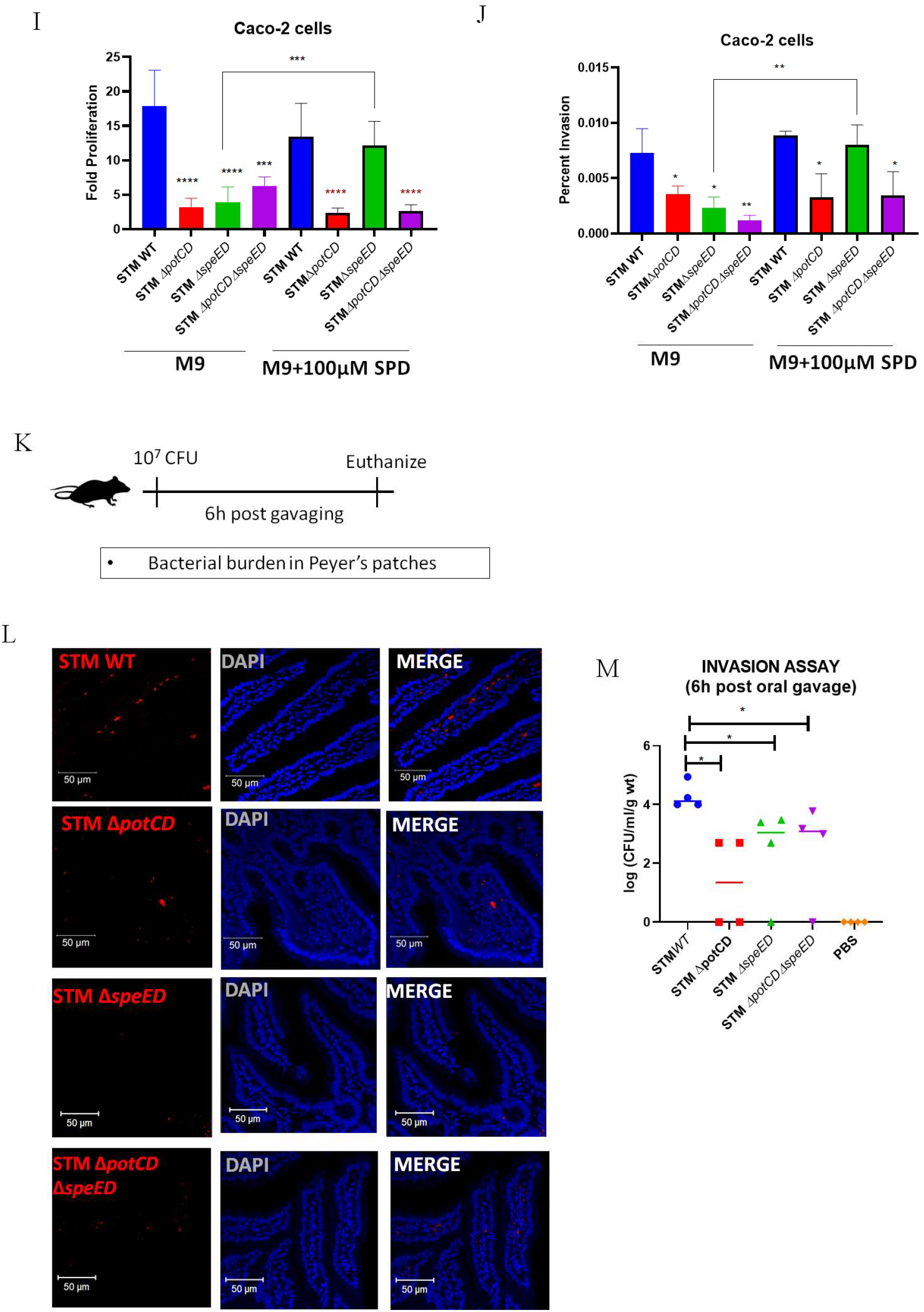
Spermidine synthesis and transport in *Salmonella* Typhimurium is essential to invade and proliferate in the host. A. The intracellular proliferation of the STM WT, STM Δ*potCD,* STM Δ*speED* and STM Δ*potCD* Δ*speED* in Caco-2 cells B. The percentage invasion into Caco-2 cells of STM WT and the mutants C. The mRNA expression of *speE* and *speD* in STM WT post infection into Caco-2 cells D. The mRNA expression of *pot-*transporter genes in STM WT post infection into Caco-2 cells E. The mRNA expression of *pot-*transporter genes in STM Δ*speED* post infection into Caco-2 cells F. The mRNA expression of *speE* and *speD* genes in STM Δ*potCD* post infection into Caco-2 cells G. The intracellular proliferation of the STM WT, STM Δ*potCD,* STM Δ*speED* and STM Δ*potCD* Δ*speED* in Caco-2 cells with supplementation of exogenous spermidine during infection H. The percentage invasion into Caco-2 cells of STM WT and the mutants with supplementation of exogenous spermidine during infection I. The intracellular proliferation of the STM WT, STM Δ*potCD,* STM Δ*speED* and STM Δ*potCD* Δ*speED* in Caco-2 cells, grown in M9 minimal media supplemented with spermidine J. The percentage invasion into Caco-2 cells of STM WT and the mutants grown in M9 minimal media supplemented with spermidine K. Experimental procedure for studying invasion of STM into mice Peyer’s patches L. Immunofluorescence of histopathological sections of mice intestine (Peyer’s patches) to study invasion, DAPI is used to stain the nucleic acids in the cells, and Anti-*Salmonella*(LPS) (Cy3 tagged secondary antibody used-Red) K. Burden of STM in Peyer’s patches post 6 hours of oral gavage to assess invasion. Student’s t-test was used to analyze the data; p values ****<0.0001, ***<0.001, **<0.01, *<0.05. Two-way Annova was used to analyze the grouped data; p values ****<0.0001, ***<0.001, **<0.01, *<0.05. For *in-vivo* studies Mann-Whitney test was used to analyze the data; p values ****<0.0001, ***<0.001, **<0.01, *<0.05.

We further assessed whether supplementation of exogenous spermidine rescue the lower fold proliferation and invasion. We supplemented the Caco-2 cells with 50µM spermidine during infection and observed that supplementation only rescued the lower fold proliferation for STM Δ*speED* (**Fig 2 G**). However, it did not rescue the reduced invasion for any of the mutants (**Fig 2 H**). STM Δ*speED* harbours the spermidine transport system, although low but may aid in the uptake of the exogenous spermidine to rescue the phenotype. However, when supplemented in minimal M9 media during the *in-vitro* growth of the different strains before infection, it rescued the lower fold proliferation and the lesser invasion of the STM Δ*speED* only (**Fig 2 I and J**). To further validate our *in*-*vitro* cell line results, we then studied the invasion of the mutants into the intestine upon orally gavaging C57BL/6 mice a sub-lethal dose of the bacteria (**Fig 2 K**). Interestingly, STM *ΔpotCD*, STM *ΔspeED*, and STM *ΔpotCD ΔspeED* invaded significantly less into the Peyer’s patches of the mice (**Fig 2 L and M**). Thus, *Salmonella* requires spermidine to invade successfully, subsequently survive, and proliferate within the epithelial cells *in-vitro*and *in-vivo*.

### Adhesion of *Salmonella* Typhimurium to epithelial cells is aided by spermidine by regulation of fimbrial and non-fimbrial adhesins

Most bacterial pathogens must first reach the site of infection, followed by a sequential steps of adhesion, invasion, multiplication and proliferation to infect and colonise the host tissues successfully [29]. During the pathogenesis of *Salmonella*, a crucial step towards infection into the IECs is its ability to adhere to the surface of the IECs [30]. Our study so far shows that the loss of spermidine transport or synthesis capability of *Salmonella* reduces the invasiveness of the bacteria into human epithelial cells. Polyamines such as cadaverine have been shown to play an important role in the pathogenesis of respiratory tract pathogens like *S. pneumoniae* by aiding in the stages of adhesion leading to colonisation in the nasopharynx [31]. Also, exogenous spermidine increased the adhesion of *Bifidobacterium animalis subs. lactis* Bb12 in the mucous of infants [32]. To understand the role of spermidine in the adhesion of *Salmonella* Typhimurium to epithelial cells, we performed an adhesion assay in Caco-2 cells with various strains. All three mutants showed significantly lower adhesion than the wild-type ones (**Fig 3 A**). This was also found in HeLa cells (**Fig S3 A**). To validate our observation we performed immunofluorescence and found reduced adhesion of the transport and biosynthesis mutants to Caco-2 cells (**Fig 3 C and D**). The addition of exogenous polyamine to the bacteria before infection reversed the phenotype in STM *ΔspeED* (**Fig 3 B, C and D**).

**Fig 3:**
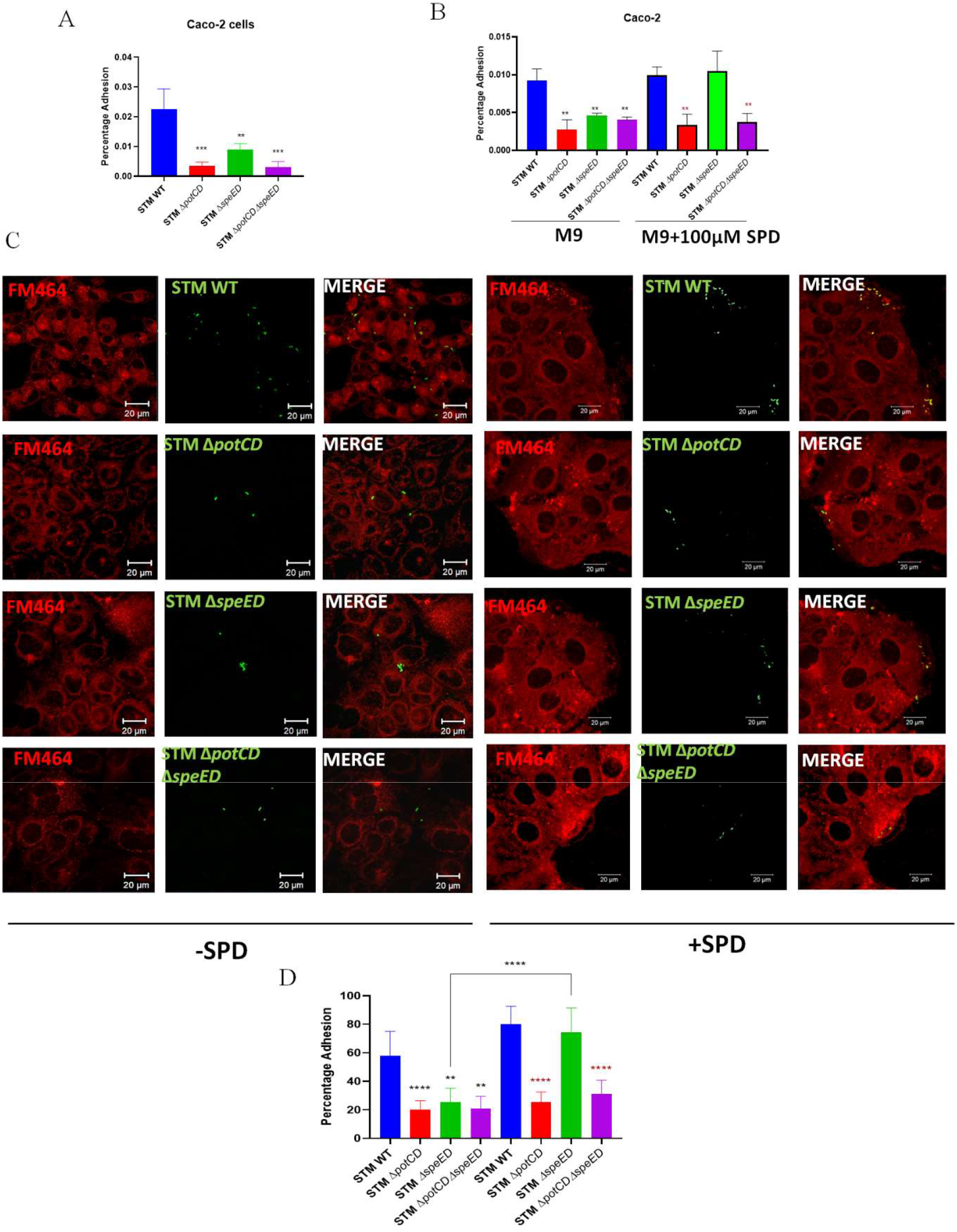
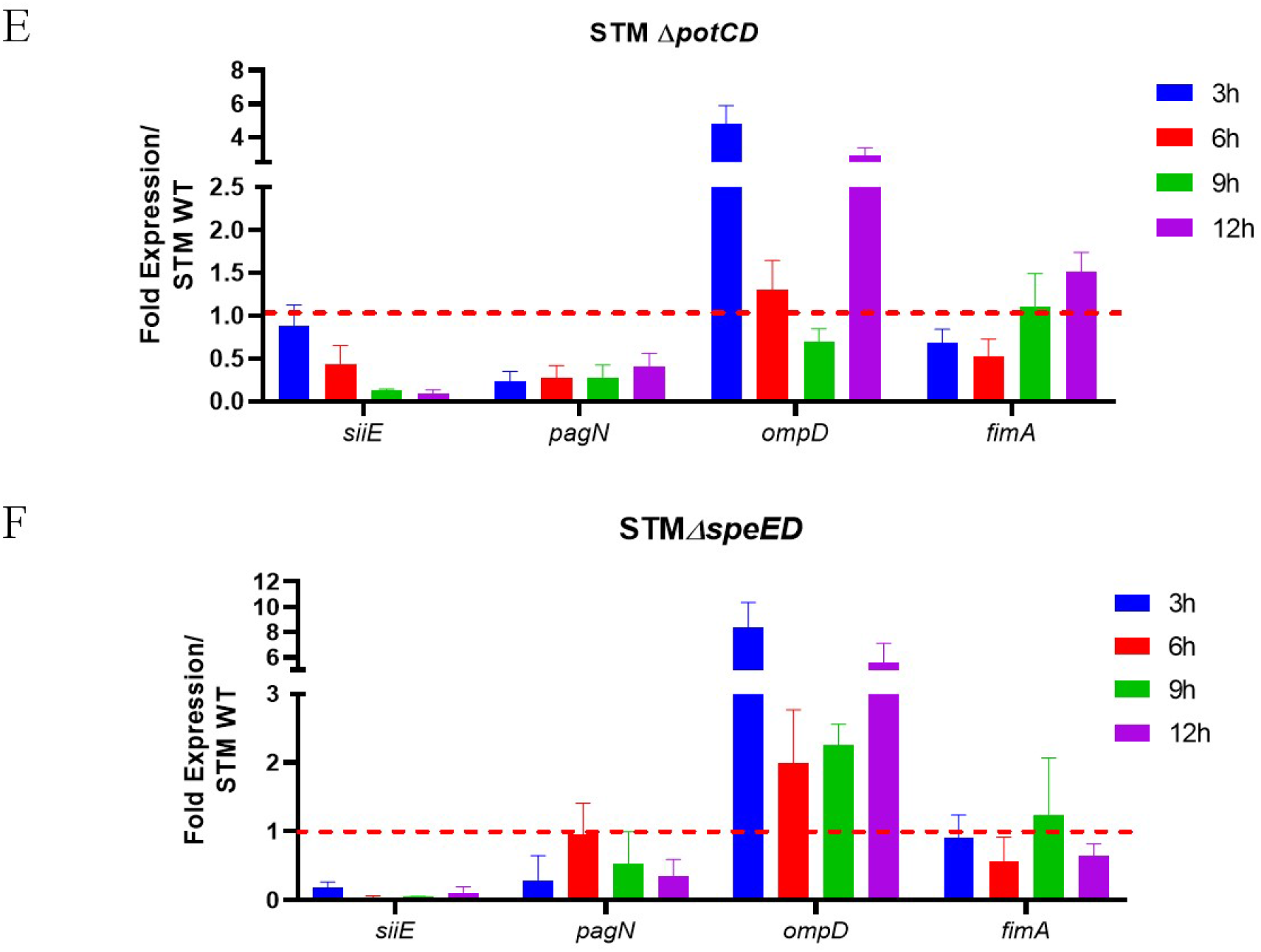
Adhesion of *Salmonella* Typhimurium to epithelial cells is aided by spermidine by regulation of fimbrial and non-fimbrial adhesins. A. Adhesion assay of STM WT and the mutants Caco-2 cells B. Adhesion assay in Caco-2 of STM WT and the mutants grown in M9 minimal media supplemented with spermidine C. Immunofluorescence imaging to study the adhesion to Caco-2 cells of STM WT and the three mutants grown in M9 minimal media with and without supplementation of spermidine, here FM464(red) is used to stain the lipids for Caco-2 and Anti-*Salmonella*(LPS) (Alexafluor-488 tagged secondary antibody used-Green) for STM D. Quantification of C E. The mRNA expression of non-fimbrial adhesins such as *siiE, pagN* and *ompD* and fimbrial adhesin *fimA* genes in STM Δ*potCD* during *in-*vitro growth in LB media F. The mRNA expression of non-fimbrial adhesins such as *siiE, pagN* and *ompD* and fimbrial adhesin *fimA* genes in STM Δ*speED* during *in-*vitro growth in LB media. Student’s t-test was used to analyze the data; p values ****<0.0001, ***<0.001, **<0.01, *<0.05.

*Salmonella* employs multiple systems ranging from monomeric structures to highly complex and giant structures to adhere to the host cells. *Salmonella* possesses multiple fimbrial gene clusters that encode fimbrial appendages to bind to the host cell surfaces [33, 34]. Apart from fimbrial proteins, its cell surface is decorated with various non-fimbrial proteins like PagN, outer membrane proteins (Omps) and the type 1 secreted giant adhesin SiiE etc. [30, 35, 36]. We observed that the fimbrial and non-fimbrial adhesins in both mutants show lower mRNA expression than in the wild type during the exponential phase of growth (**Fig 3 E and F**). Furthermore, growth in supplementation of exogenous spermidine showed an increase in the mRNA expression in the mid-log growth phase for the non-fimbrial *siiE* and *pagN* and the fimbrial *fimA* genes (**Fig 3 G**). These observations confirm that spermidine aids in the adhesion of *Salmonella* Typhimurium to host cells by regulating the expression of fimbrial and non-fimbrial genes.

### Spermidine regulates flagellar gene expression by enhancing the translation of FliA, which otherwise has a poor Shine-Dalgarno sequence and an unusual START codon

Gram-negative and Gram-positive bacteria express flagella on their surfaces, primarily as a motility structure [37]. However, many studies have shown flagella to act as an appendage to adhere to host cell surfaces, such as the chromosomal mutation of the *fliD* gene that encodes the flagellar cap protein in *Pseudomonas aeruginosa*, which made the bacteria nonadhesive to mucin on epithelial cells [38]. Similarly, in *Vibrio cholerae,* non-motile variants exhibited reduced virulence due to poor adsorption onto the cells [39]. Likewise, in *Salmonella,* the importance of flagella as an adhesive structure has been shown by many researchers [40, 41]. Researchers from our group have previously demonstrated that the loss of flagella in *Salmonella* Typhimurium led to reduced adhesion to Caco-2 cells and lesser colonisation in the gut of *C. elegans* [20]. Furthermore, the global transcriptomic analysis in the *speG* deletion mutant of *Salmonella* Typhimurium, showed that the genes associated with the regulation and formation of the flagella were downregulated [18]. Thus, we were interested in deciphering the role of spermidine in regulating flagella. We carried out a swimming motility assay and observed that STM *ΔpotCD* and STM *ΔpotCDΔspeED* showed highly attenuated movement on soft agar (**Fig 4 A, S4 A**). In contrast, STM *ΔspeED* exhibited a 40 percent reduction in the movement than the wild type (**Fig 4 A, S4 A**). Thus, we determined the mRNA expression of the *fliC and fljB* that encode flagellin protein in *Salmonella* Typhimurium and found that both the genes show significant downregulation in STM *ΔpotCD* and STM *ΔspeED* (**Fig 4 B and C**). However, the growth of STM *ΔspeED* in the presence of spermidine increased the swimming motility similar to wild type and the mRNA expressions of *fliC* and *fljB* during the mid-log phase of growth (**Fig 4 A and D**). This was not observed for STM *ΔpotCD*.

**Fig 4:**
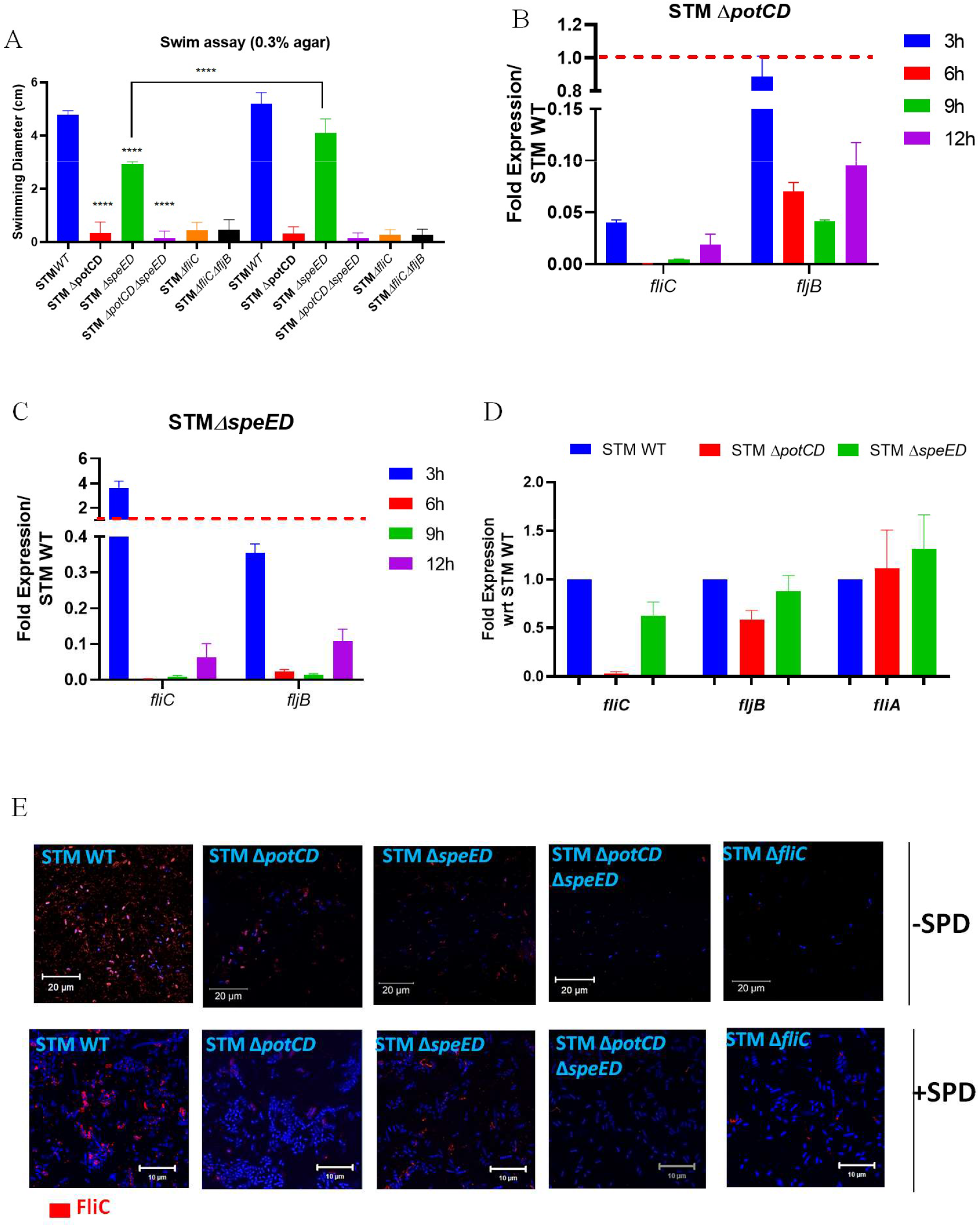
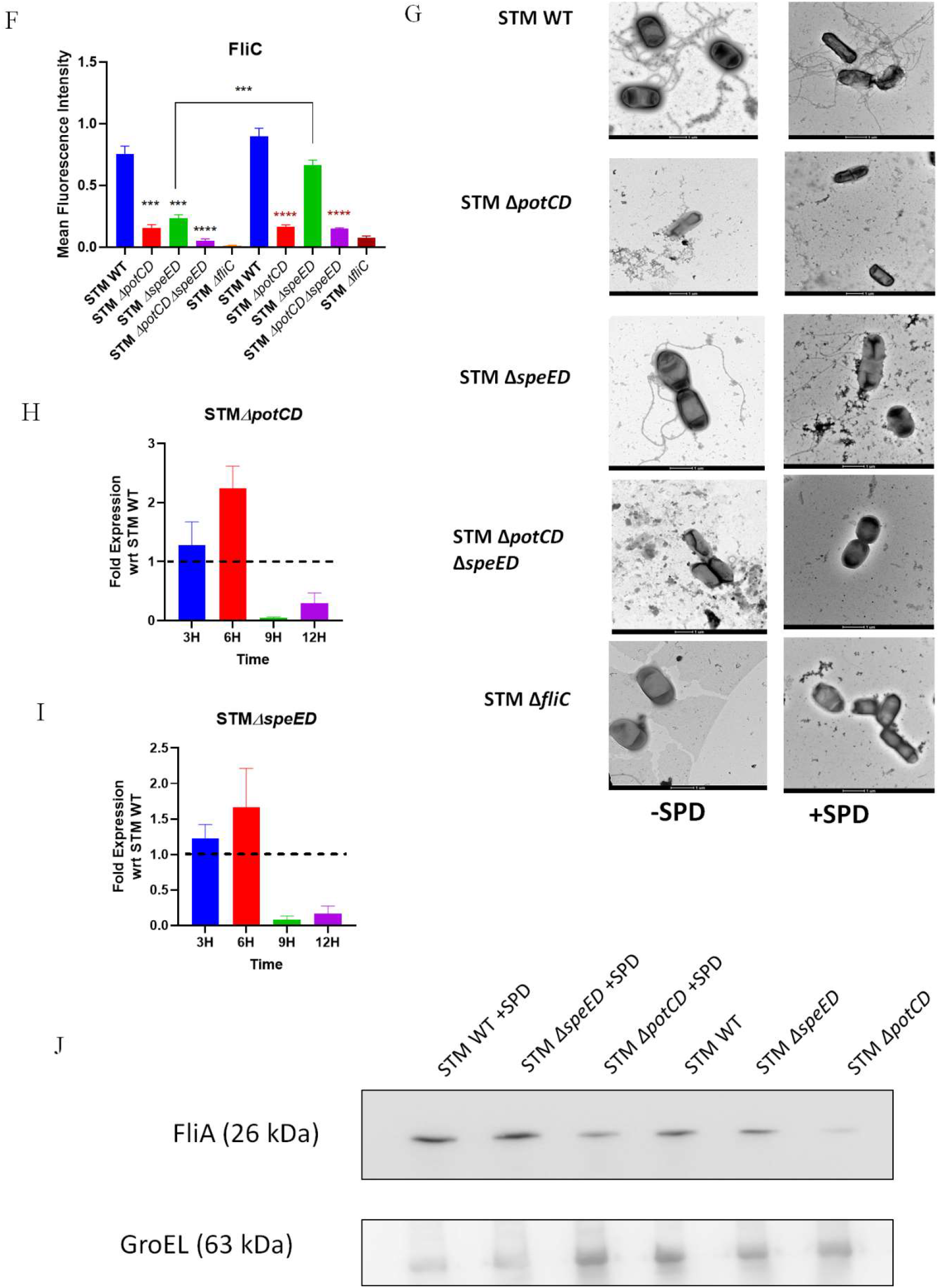
Spermidine regulates flagellar gene expression by enhancing the translation of FliA, which otherwise has a poor Shine-Dalgarno sequence and an unusual START codon. A. Swimming motility of STM WT, STM Δ*potCD,* STM Δ*speED* and STM Δ*potCD* Δ*speED* grown in M9 minimal media supplemented with and without spermidine on 0.3% agar, B. The mRNA expression of the genes *fliC* and *fljB,* coding for flagellin in STM Δ*potCD* during *in-* vitro growth in LB media, C. The mRNA expression of the genes *fliC* and *fljB,* coding for flagellin in STM Δ*speED* during *in-*vitro growth in LB media, D. The mRNA expression of the genes *fliC* and *fljB* coding for flagellin and *fliA*, coding for the sigma-factor-28 in STM Δ*potCD* and STM Δ*speED* during log phase of growth (6 hours) in LB media supplemented with spermidine, E. Immunofluorescence imaging to study the expression of FliC (flagella) on the surface of STM WT and the three mutants grown with and without supplementation of spermidine, here DAPI (blue) is used to stain the nucleoid of STM and Anti-FliC(Cy3-tagged secondary antibody used-Red) for STM, F. The quantification of E, G. TEM images of STM WT, STM Δ*potCD,* STM Δ*speED*, STM Δ*potCD* Δ*speED* and STM Δ*fliC* grown till log phase of growth,with and without supplementation of spermidine, H. The mRNA expression of the gene *fliA,* coding for sigma-factor-28 in STM Δ*potCD* during *in-*vitro growth in LB media, I. The mRNA expression of the gene *fliA,* coding for sigma-factor-28in STM Δ*speED* during *in-* vitro growth in LB media, J. Western blot of *fliA*-FLAG in STM WT, STM Δ*potCD* and STM Δ*speED* grown till log phase of growth,with and without supplementation of spermidine. Student’s t-test was used to analyze the data; p values ****<0.0001, ***<0.001, **<0.01, *<0.05. Two-way Annova was used to analyze the grouped data; p values ****<0.0001, ***<0.001, **<0.01, *<0.05.

We performed immunofluorescence to study the flagellin FliC expression on *Salmonella* Typhimurium’s surface. The presentation of the flagellin FliC was significantly less on the surface in all the mutants than in the wild type (**Fig 4 E and F, S4 B**). The result was further validated using Transmission electron microscopy. Likewise, we observed reduced numbers of flagella on the surface of STM *ΔspeED*, while no flagella on the surface of STM *ΔpotCD* and STM *ΔpotCDΔspeED* similar to STM *ΔfliC* (**Fig 4 G**). The surface presentation of FliC was found to be more upon growth of STM *ΔspeED* in the presence of exogenous spermidine (**Fig 4 E-G)**. As the levels of *fliC* and *fljB* decreased in both the spermidine transport and biosynthesis mutants of *Salmonella*, we next determined the expression of the sigma factor σ^28^ (FliA) that aids in the transcription of the flagellin genes. In both STM *ΔpotCD* and STM *ΔspeED*, the mRNA expression was not downregulated in the exponential to mid-log growth phase (**Fig 4 H and I**). Polyamines, cationic molecules, interact with the negatively charged nucleic acids and often regulate the transcription and translation of multiple genes. The genes that are regulated by polyamines fall under the polyamine regulon. Multiple sigma factors such as *rpoS*, *hns, oppA* etc., are known to be under the polyamine modulon [42, 43]. In *E. coli* OppA has a weak and distant Shine-Dalgarno sequence and polyamines stimulate the translation of OppA in such a case [44, 45]. Also, polyamines increase the translation of RpoN and H-NS whose transcript contains a poor Shine-Dalgarno (SD) sequence in *E. coli* and that of Cra which possesses an unusual “GUG” start codon in its transcript [46]. Interestingly, in *Salmonella* Typhimurium *fliA,* the transcript likewise contains an unusual “GTG” START codon and a poor SD sequence located farther than 6-7bps from the START codon **(S4 C**). Thus, we tagged *fliA* with FLAG-Tag in the chromosome of STM WT, STM *ΔpotCD* and STM *ΔspeED,* and observed that there was significant downregulation of FliA in both STM *ΔpotCD* and STM *ΔspeED*. There was an increase in the expression of FliA when STM *ΔspeED* was grown in the presence of exogenous polyamine (**Fig 4 I, S4 D**). These results show that spermidine regulates the expression of flagellin genes by enhancing the translation of FliA (σ^28^), which otherwise possess an unusual START codon and a poor SD sequence in *Salmonella* Typhimurium.

### Spermidine tunes the expression of the *Salmonella* pathogenicity island-1, thereby facilitating the invasion into epithelial cells

*Salmonella* employs multiple ways to invade the IECs, of which an effective strategy is to induce its uptake by the otherwise non-phagocytic cells. *Salmonella* pathogenicity island-1 (SPI-1) encodes an elaborate nano injection machinery, the type-3 secretion system (T3SS) and innumerable effector proteins that are involved in inducing the uptake by epithelial cells [47]. The initial attachment of the bacteria to the mucin and the cell surface activates a complex intracellular regulatory network leading to the formation of the T3SS on the surface that penetrates the host cell membrane and translocates multiple effectors into the host cytosol [7, 48]. Multiple environmental signals such as osmolarity, pH, and oxygen concentration activate the SPI-1 through the master regulator HilA. Apart from these bile acids, short-chain fatty acids and magnesium ion concentration also stimulate the expression of SPI-1 genes in *Salmonella* [49]. A research group has previously shown that polyamines regulate the SPI-1 and the translation of *hilA* in *Salmonella* Typhimurium [12, 50]. This motivated us to understand the role of spermidine in regulating of *Salmonella* pathogenicity island-1. We determined the expression of SPI-1 genes in STM WT, STM *ΔpotCD* and STM *ΔspeED* during their *in-vitro* growth in LB media and observed that all the genes were significantly downregulated in both the mutants (**Fig 5 A and B**). We further validated the results by using *lacZ* constructs under promoter of *hilA* and *spiC.* The LacZ activity was significantly low in STM *ΔpotCD* and STM *ΔspeED* when cloned under the *hilA* promoter. Upon growth of STM *ΔspeED* with supplementation of spermidine, the LacZ activity was high (**Fig S5 A and B)**. This suggests that deleting of spermidine transporter and biosynthesis genes in *Salmonella* Typhimurium reduces *hilA* transcription. On the contrary, we did not observe a difference in LacZ activity when cloned under *spiC* promoter (**Fig S5 C**). The mRNA expression of the SPI-1 genes was also reduced in both STM *ΔpotCD* and STM *ΔspeED* post-infection into Caco-2 cells (**Fig 5 C and D**). As the SPI-1 genes have a highly complex regulatory network, we accessed the mRNA expression of the essential two-component system BarA/SirA, upstream of *hilA* in the regulatory network. Both *barA* and *sirA* mRNA levels were significantly low in STM *ΔpotCD* and STM *ΔspeED* than the wild type (**Fig 5 E and F**).

**Fig 5:**
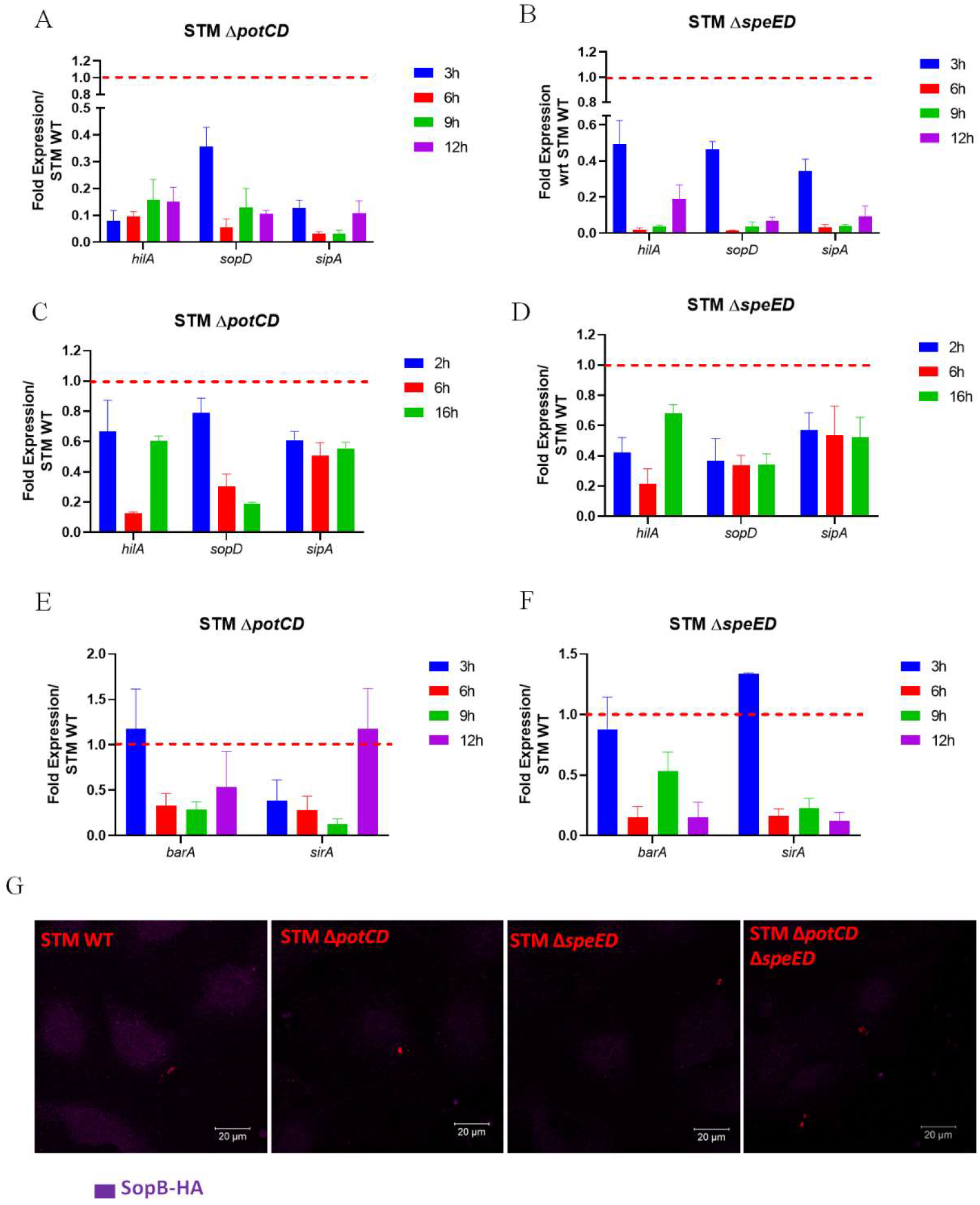
Spermidine tunes the expression of the *Salmonella* pathogenicity island-1, thereby facilitating the invasion into epithelial cells. A. The mRNA expression SPI-1 master-regulator and effectors such as *hilA, sopD* and *sipA* respectively in STM Δ*potCD* during *in-*vitro growth in LB media, B. The mRNA expression of the SPI-1 master-regulator and effectors such as *hilA, sopD* and *sipA* respectively in STM Δ*speED* during *in-*vitro growth in LB media, C The mRNA expression of SPI-1 master-regulator and effectors such as *hilA, sopD* and *sipA* respectively in STM Δ*potCD* post infection into Caco-2 cells D. The mRNA expression SPI-1 master-regulator and effectors such as *hilA, sopD* and *sipA* respectively in STM Δ*speED* post infection into Caco-2 cells, E. The mRNA expression the two-component system *barA* and *sirA*, that regulates the SPI-4 and SPI-1, in STM Δ*potCD* during *in-* vitro growth in LB media, F. The mRNA expression the two-component system *barA* and *sirA*, that regulates the SPI-4 and SPI-1 in STM Δ*speED* during *in-*vitro growth in LB media, G. Immunofluorescence imaging to study the localisation of SopB-HA (Anti-HA, secondary antibody tagged with Alexafluor-488, pseudo-coloured to magenta) in STM WT and the three mutants (expressing RFP-red), 2 hours post infection into Caco-2 cells. Student’s t-test was used to analyze the data; p values ****<0.0001, ***<0.001, **<0.01, *<0.05. Two-way Annova was used to analyze the grouped data; p values ****<0.0001, ***<0.001, **<0.01, *<0.05.

Along with SPI-1 encoded effectors, the T3SS of SPI-1 also translocate an SPI-5 encoded effector protein SopB. SopB is a phosphoinositide-phosphatase, and researchers from our group have previously shown the role of this SPI-1 effector in *Salmonella* Typhimurium virulence [51]. We used HA-tagged SopB to understand the regulation of SPI-1 in STM *ΔpotCD* and STM *ΔspeED*. Using immunofluorescence, we noticed that, indeed in the two mutants, the translocation of SopB is lesser than in the wild type (**Fig 5 G**). Our results thus explain the crucial role of spermidine.

## Discussion

The pathogenesis of most pathogens involves the entry into the host tissues and cells. However, to enter the host cells, the pathogens require multiple crucial steps before entering the host. These include motility to reach the site of entry, the initial attachment and adhesion to the cell surfaces and subsequent invasion using multiple strategies. *Helicobacter pylori* was initially considered a non-invasive pathogen; however, studies in the past decade have proven it invasive. To infect *Helicobacter pylori* initially attaches to the cell surface, followed by receptor-mediated invasion and survival [52]. The Gram-positive pathogen *Staphylococcus aureus* expresses fibronectin-binding proteins that help them adhere to the eukaryotic cells, promoting uptake by the host cells [53]. *Salmonella* is an enteric pathogen which enters the host system through the feco-oral route, and at the intestine, it crosses the intestinal barrier to cause systemic infection. Likewise, during *Salmonella* pathogenesis, it must first attach to the mucin in the intestinal lumen and to the cell surface to gain entry into the intestinal epithelial cells [54]. *Salmonella* is considered an ancient pathogen associated with the death of humankind for thousands of years [55, 56]. Over the years, it has emerged as a successful enteric pathogen by modulating its strategies and employing diverse arms and shields, allowing it to conquer diverse niches during its pathogenesis. The complexity of the arsenals and the surface structures forces us to develop combating strategies against the disease-causing pathogen.

Polyamines are ubiquitously present in all living forms, including prokaryotes. In eukaryotes, polyamines are often crucial for cellular functions and, most importantly, cell division. Thus, tumour cells are the prime site expressing high levels of polyamines to cope with the high rate of cell division. In prokaryotes, they are linked to the growth, and stress response by regulation of the expression of multiple genes. Few research groups have shown that polyamines are essential in *Salmonella’s* virulence and stress response [12]. However, a dearth of mechanistic understanding remains in the field. Of the various polyamines, putrescine, spermidine and cadaverine are the major ones in *Salmonella*. Most of the studies show the role of the group of polyamines, rather than any particular polyamines, in the virulence of *Salmonella*. Thus, we were fascinated to delve into the mechanism behind the role of the vital polyamine, spermidine, in *Salmonella* pathogenesis. Also, *Salmonella* imports as well as synthesises intracellularly, it is intriguing to understand the regulation of the two processes.

For the first time, we report that the two processes are not mutually exclusive rather, *Salmonella* interestingly regulates them together. It suggests that the transporter and the synthesis genes are required to function together to maintain the intracellular homeostasis of spermidine. The *speG* is vital in removing the accumulated spermidine, however our study shows that *Salmonella* has more complexity in maintaining the homeostasis of spermidine intracellularly. *Salmonella* utilises the small molecules like spermidine to regulate the expression of multiple adhesive and non-adhesive complex surface structures, thus identifying new members of polyamine regulon in *Salmonella*. Spermidine also regulates the elegant nano-injection machinery of the T3SS and the effectors of the SPI-1 for its uptake and survival host cells. We expect that spermidine binds to the anionic nucleic acids and there by tune the expression of the multiple genes in *Salmonella.* However, further study is essential in unravelling the mechanism of interaction of spermidine to the nucleic acid in *Salmonella.* We report a novel regulatory pathway in *Salmonella*, where we show that spermidine aid in overcoming the obstacle of a weak and poor transcript, thereby maintaining the synthesis of the elaborate surface tools required for motility and attachment. As previously explained that the pathogenic bacteria use a complex network of molecules to evade and survive. Our study solves the enigma of how spermidine regulates diverse aspects and essentially is a novel player in the complex network regulating the virulence in *Salmonella.* This study opens avenues to design drugs that target the polyamine metabolism in *Salmonella* and thus reducing the infectivity and the burden of *Salmonella*.

## Acknowledgement

Prof. Umesh Varshney and Mr. Jitendra Bisht from MCB, IISc are duly acknowledged for providing the plasmid for FLAG-tag generation. Departmental Confocal Facility, Departmental Real-Time PCR Facility, Divisional MS facility, Divisional EM facility and Central Animal Facility at IISc are duly acknowledged. Mr Sumith and Ms Navya are acknowledged for their help in image acquisition. Mrs. Sunita is duly acknowledged for helping with mass spectrometry. Mr. Prakhar is also acknowledged for technical help.

## Author contribution statement

AVN and DC conceived the study. AVN and DC designed experiments. AVN performed experiments. AVN, YD and SAR performed LC QTOF MS/MS experiment, and UST provided valuable inputs for the LC QTOF MS/MS experiment. AVN, analysed the data, prepared the figures and wrote the manuscript draft. AVN and DC reviewed and edited the manuscript. DC supervised the work. All the authors read and approved the manuscript.

