## supplementary for "Spermidine facilitates the adhesion and subsequent invasion of *Salmonella* Typhimurium into epithelial cells via the regulation of surface adhesive structures and the SPI-1"

Dipshikha Chakravortty

Email: dipa@ iisc.ac.in

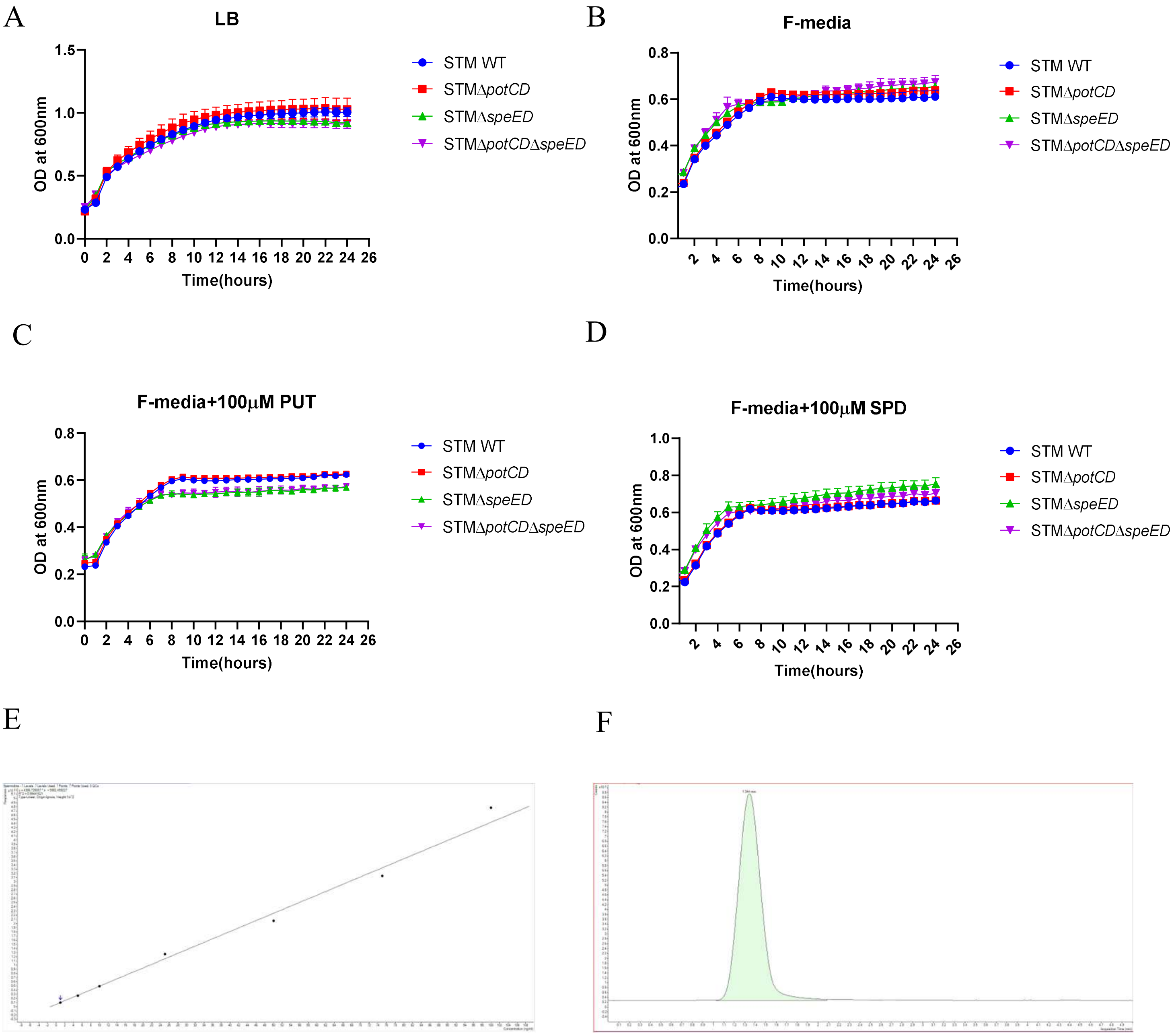


**S1.** A. The growth kinetics of STM WT, STM Δ*potCD*, STM Δ*speED* and STM Δ*potCD*Δ*speED* in LB media, B. The growth kinetics of STM WT, STM Δ*potCD*, STM Δ*speED* and STM Δ*potCD*Δ*speED* in acidic F-media that mimics SCV, C. The growth kinetics of STM WT, STM Δ*potCD*, STM Δ*speED* and STM Δ*potCD*Δ*speED* in acidic F-media that mimics SCV, supplemented with Putrescine (PUT), D. The growth kinetics of STM WT, STM Δ*potCD*, STM Δ*speED* and STM Δ*potCD*Δ*speED* in acidic F-media that mimics SCV, supplemented with Spermidine (SPD), E. Standard curve with pure SPD using LC-MS/MS, F. Chromatogram for SPD detection using LC-MS/MS.

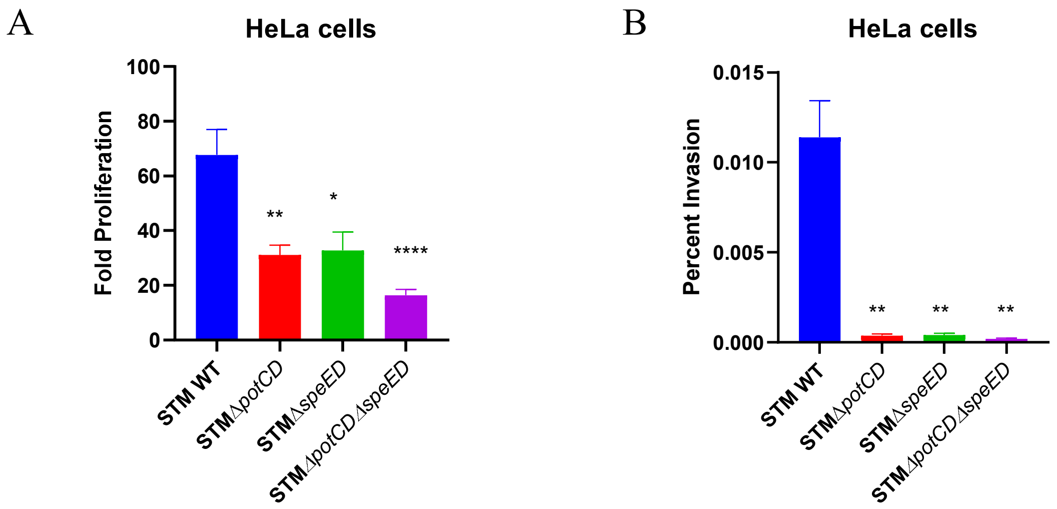


**S2.** A. The intracellular proliferation of the STM WT, STM ΔpotCD, STM ΔspeED and STM ΔpotCD ΔspeED in HeLa cells B. The percentage invasion into HeLa cells of STM WT and the mutants

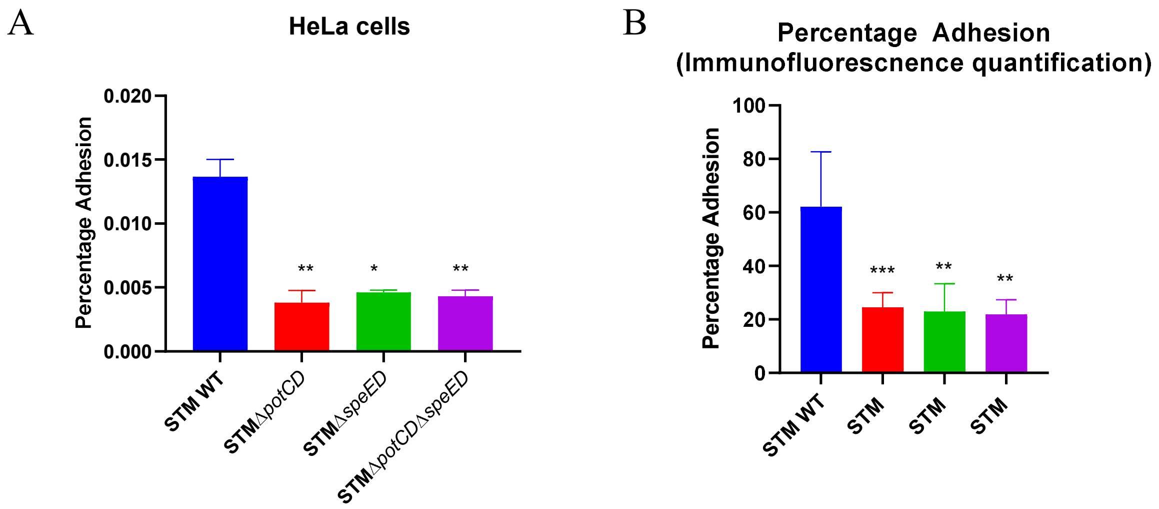


**S3.** A. Adhesion assay of the STM WT, STM Δ*potCD*, STM Δ*speED* and STM Δ*potCD*Δ*speED* in HeLa cells B. Quantification of same using immunofluorescence.

A

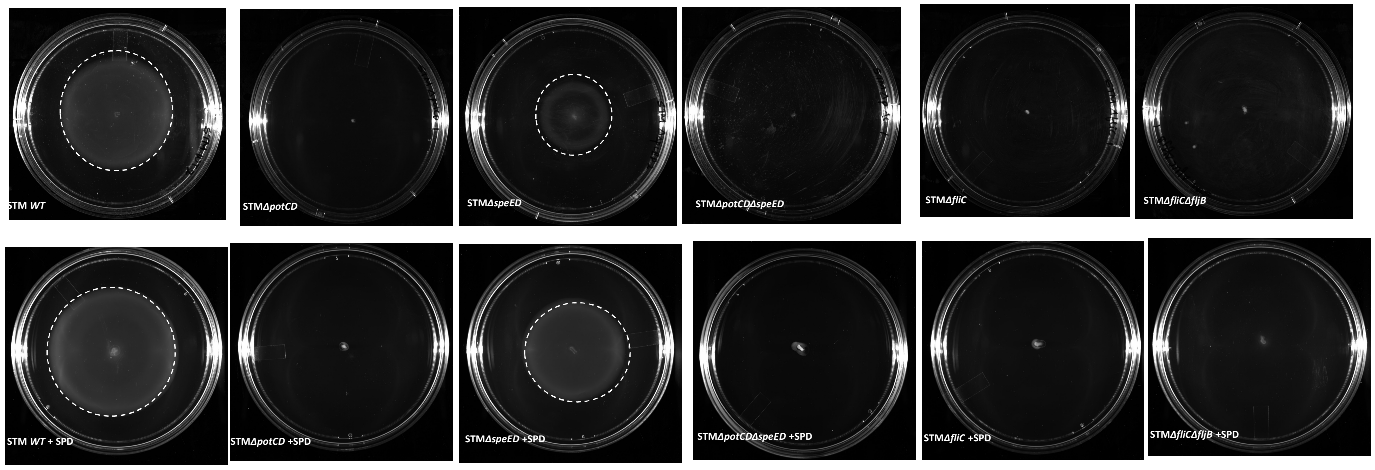


B

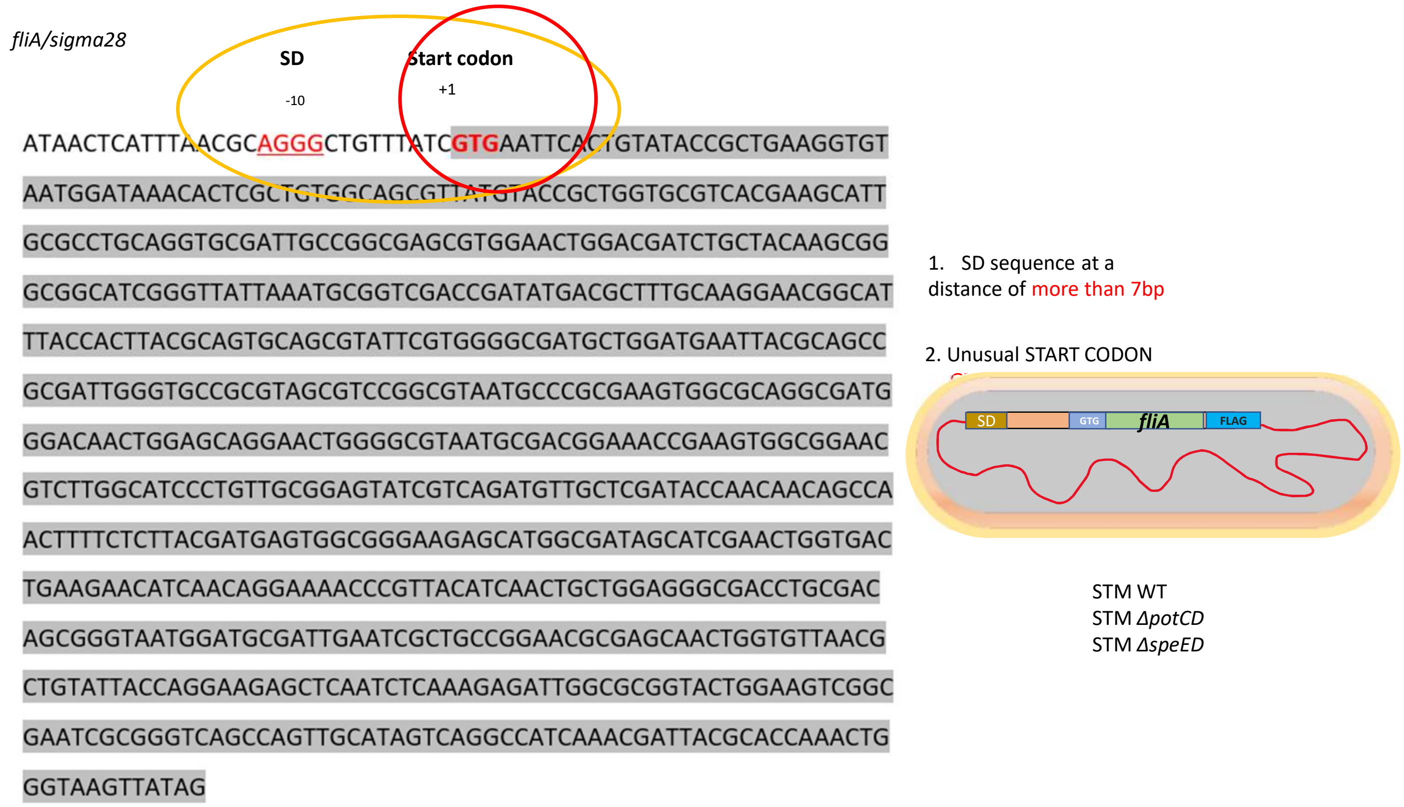


C

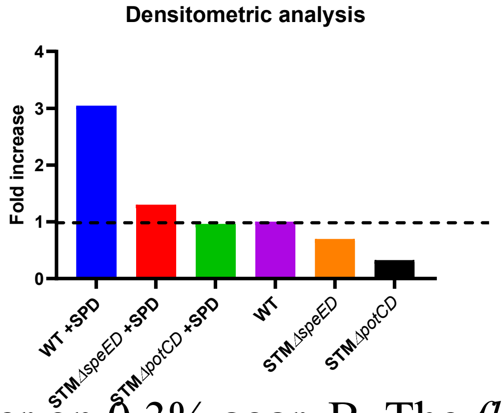


**S4.** A. Swimming diameter on 0.3% agar, B. The *fliA* having an unusual START codon and a poor SD sequence, and the strategy to chromosomally tag *fliA* eith FLAG in STM WT, STM Δ*potCD*, STM Δ*speED,* C desitometric plot for immunoblot( representative image of one-time experiment. This experiment was repeated atleast thrice to validate our results.

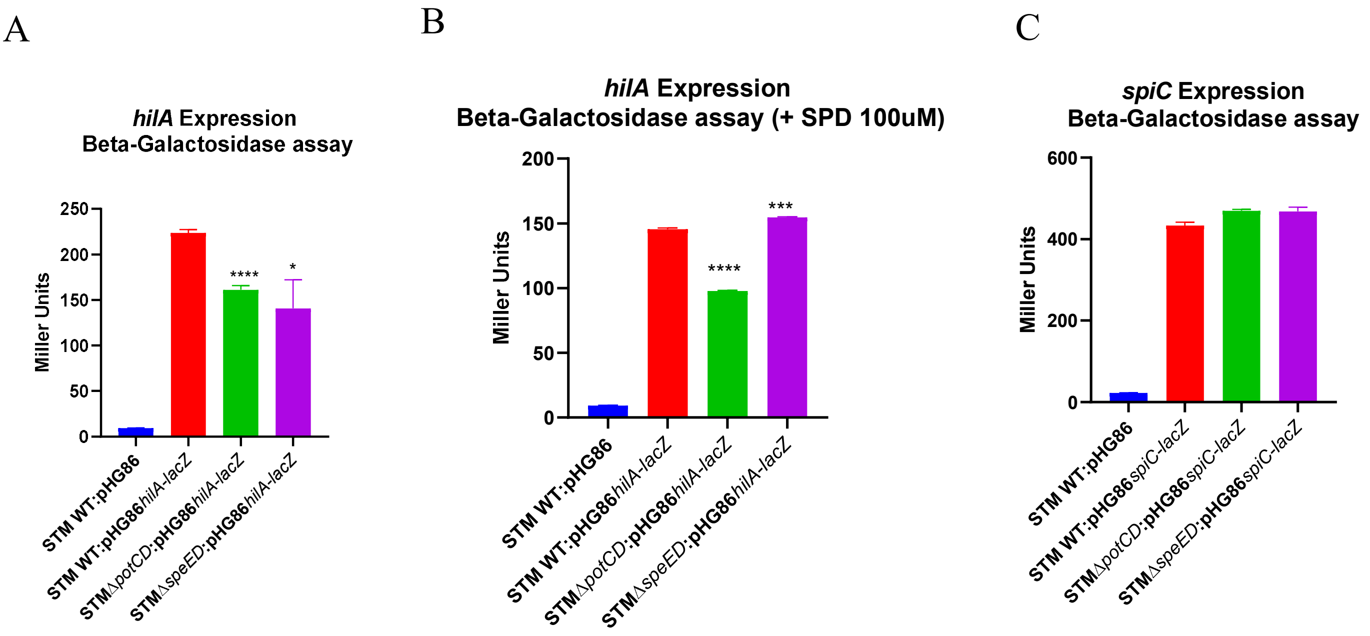


D

Miller Unit (MU)= 1000[ $OD_{420nm}$ $OD_{550nm}$ * 1.75]/ T* V* $OD_{600nm}$

$OD_{550nm}$ – Light scattering by cell debris
$OD_{600nm}$ – Bacterial cell density
$OD_{420nm}$ – Absorbance by o-nitrophenol and light scattering by cell debris
T- Time of the reaction in minutes
V- Volume of the culture in mL

**S5.** A. Lac-Z assay in STM WT, STM Δ*potCD*, STM Δ*speED* expressing *lacZ* under *hilA* promoter upon growth without spermidine, B. growth with supplementation of exogenous spermidine,C. *lacZ* cloned under *spiC* promoter, D. Miller Unit calculation for determination of *hilA* and *spiC* promoter activity.
